# Maize GO Annotation - Methods, Evaluation, and Review (maize-GAMER)

**DOI:** 10.1101/222836

**Authors:** Kokulapalan Wimalanathan, Iddo Friedberg, Carson M. Andorf, Carolyn J. Lawrence-Dill

## Abstract

We created a new high-coverage, robust, and reproducible functional annotation of maize protein coding genes based on Gene Ontology (GO) term assignments. Whereas the existing Phytozome and Gramene maize GO annotation sets only cover 41% and 56% of maize protein coding genes, respectively, this study provides annotations for 100% of the genes. We also compared the quality of our newly-derived annotations with the existing Gramene and Phytozome functional annotation sets by comparing all three to a manually annotated gold standard set of 1,619 genes where annotations were primarily inferred from direct assay or mutant phenotype. Evaluations based on the gold standard indicate that our new annotation set is measurably more accurate than those from Phytozome and Gramene. To derive this new high-coverage, high-confidence annotation set we used sequence-similarity and protein-domain-presence methods as well as mixed-method pipelines that developed for the Critical Assessment of Function Annotation (CAFA) challenge. Our project to improve maize annotations is called maize-GAMER (GO Annotation Method, Evaluation, and Review) and the newly-derived annotations are accessible via MaizeGDB (http://download.maizegdb.org/maize-GAMER) and CyVerse (B73 RefGen_v3 5b+ at doi: doi.org/10.7946/P2S62P and B73 RefGen_v4 Zm00001d.2 at doi: doi.org/10.7946/P2M925).

## 2 Introduction

Maize is an agriculturally important crop species and model organism for genetics and genomics research (Lawrence et al., 2004). Not only is maize historically important for genetics research, along with other model species, significant efforts have been made to transition existing datasets into a more sequence-centric paradigm (Sen et al., 2009), thus enabling genomics approaches to be brought to bear on both basic research problems and applied breeding (Lawrence et al., 2008). In 2009 the maize genome’s reference sequence was made available to the research community (Schnable et al., 2009). Since then, much work has gone into improving the utility of the genome sequence to scientists with a focus on sequence annotation.

In practice, making a genome sequence useful involves three basic steps: assembling the genome sequence, assigning gene structures, and assigning functions to genes. The quality of data generated at each step influences downstream inferences, with high-quality sequence, assembly, and gene structure assignments generally resulting in better functional annotations overall. Functional predictions serve as the basis for formulating hypotheses that are subsequently tested in the lab. As such, experimentalists have a great interest in high-quality functional annotation sets that cover all or most of the genes in their species of interest.

The Gene Ontology (GO) is a controlled vocabulary of hierarchically related terms that describe gene product function. It consists three categories: Biological Process (BP), Cellular Component (CC), and Molecular Function (MF). In the context of GO, functional annotation of a gene consists of the assignment of one or more GO terms from one or more of the GO categories to a given gene or gene model (here we will refer to genes and gene models simply as ‘genes’ for simplicity).

For individual GO term associations to genes, Evidence Codes (ECs) are assigned to assert how the association of term to gene was made (Harris et al., 2004). GO evidence codes are aggregated into five general categories: Experimental, Computational Analysis, Curator Statement, Author Statement, and Automatically Assigned. See Table 1 and http://www.geneontology.org/page/guide-go-evidence-codes for a detailed explanation of the GO evidence codes.

**Table 1:** Evidence Codes used in gene ontology annotations

| Type | Evidence Code | Description |
| --- | --- | --- |
| Experimental | EXP | Inferred from Experiment |
|  | IDA | Inferred from Direct Assay |
|  | IPI | Inferred from Physical Interaction |
|  | IMP | Inferred from Mutant Phenotype |
|  | IGI | Inferred from Genetic Interaction |
|  | IEP | Inferred from Expression Pattern |
| Computational Analysis | ISS | Inferred from Sequence or structural Similarity |
|  | ISO | Inferred from Sequence Orthology |
|  | ISA | Inferred from Sequence Alignment |
|  | ISM | Inferred from Sequence Model |
|  | IGC | Inferred from Genomic Context |
|  | IBA | Inferred from Biological aspect of Ancestor |
|  | IBD | Inferred from Biological aspect of Descendant |
|  | IKR | Inferred from Key Residues |
|  | IRD | Inferred from Rapid Divergence |
|  | RCA | Inferred from Reviewed Computational Analysis |
| Author statement | TAS | Traceable Author Statement |
|  | NAS* | Non-traceable Author Statement |
| Curatorial Statement | IC | Inferred by Curator |
|  | ND* | No biological Data available |
| Automatically-Assigned | IEA* | Inferred from Electronic Annotation |

\* Indicates evidence codes without either curation or biological data supporting them.

The use of experimental ECs asserts that the assignment results from a physical characterization of the protein’s function as described in a publication. Computational approaches are based on *in silico* analyses. One of the simplest and most commonly conducted computational approaches involves matching similar genes between an existing, well-annotated genome and an unannotated genome. Once the matches are assigned, annotations are inferred to genes in the unannotated genome. Such assignments receive the ISS (Inferred from Sequence or Structural Similarity) EC. The ISS EC is also assigned if an uncharacterized sequence contains a characterized domain. In such instances, the presence of the domain itself can be used to predict function for the uncharacterized sequence. For Curator and Author Statements, included EC types are based on judgment by curators and scientists in their expert opinion. As such, they are considered to be reviewed annotation types, though these do include two ECs based on little data: NAS (Non-traceable Author Statement) and ND (No biological Data available). The Automatically Assigned EC type contains only one EC: Inferred from Electronic Annotation (IEA). IEA is unique in that no reviewed analysis of the assignment is required. Put another way, no curatorial judgment is applied, making it the least supported EC of the group.

Sequence-based approaches to automated functional annotation generally fall into three basic categories: sequence-similarity, domain-based methods, and mixed-methods. Sequence-similarity based gene matching most often relies on BLAST (e.g., BLAST2GO) followed by limiting the number of accepted matches based on e-value or a reciprocal-best-hit (RBH) strategy (Altschul et al., 1990; Conesa and Götz, 2008; Moreno-Hagelsieb and Latimer, 2008). Domain-based methods score sequences for the presence of well-described protein domain such as those included in Pfam, PANTHER, and ProSite (Finn et al., 2017). InterProScan is a commonly used domain-based GO annotation pipeline (Jones et al., 2014). Mixed-methods combine sequence-similarity, domain-based approaches, and other evidence such as inferred orthology through phylogenetics to assign GO terms systematically (Clark and Radivojac, 2011; Falda et al., 2012; Koskinen et al., 2015). For more of the latest methods, see (Jiang et al., 2016).

For maize, two genome-scale GO annotation sets exist for the B73 reference assembly and gene set (i.e., B73 RefGen_v3 and 5b+, respectively). These functional annotations are generated by and accessible from the Gramene (www.gramene.org; (Tello-Ruiz et al., 2016)) and Phytozome (phytozome.jgi.doe.gov; (Goodstein et al., 2012)) projects and websites, respectively. Gramene annotations are based on the Ensembl annotation pipeline (http://ensemblgenomes.org/info/data/cross_references), which is a mixed-method approach. The primary sources of the Ensembl annotations are from UniProtKB, community-based annotations from MaizeGDB (Andorf et al., 2016), InterPro2GO, and projections from orthologs inferred from phylogenetic analyses. Phytozome has a two-step process for GO annotation. First, Pfam domains are assigned to proteins. Second, GO annotations are determined based on the Pfam2GO mapping (Hunter et al., 2009).

Given the wealth of functional descriptions derived from mutational analyses, many researchers rely on the available maize GO-based functional annotations from large-scale, high-profile community resources like Gramene and Phytozome for formulating experimental hypotheses, and also as input datasets to transitively annotate predicted functions to newly sequenced grass species and crop genomes (e.g., (Hirsch et al., 2016)). However, if we compare the EC types for GO assignments between the model species *Arabidopsis thaliana* and the Gramene and Phytozome functional annotations of the maize reference line B73, it is clear that the evidence supporting GO term assignments for these maize datasets is comparatively lacking (see Figure 1). Both the Gramene and Phytozome maize annotations have few annotations beyond those Inferred from Electronic Annotation (IEA). This situation is not intuitive to researchers given that maize has a wealth of functional descriptions in the literature.

**Figure 1:**
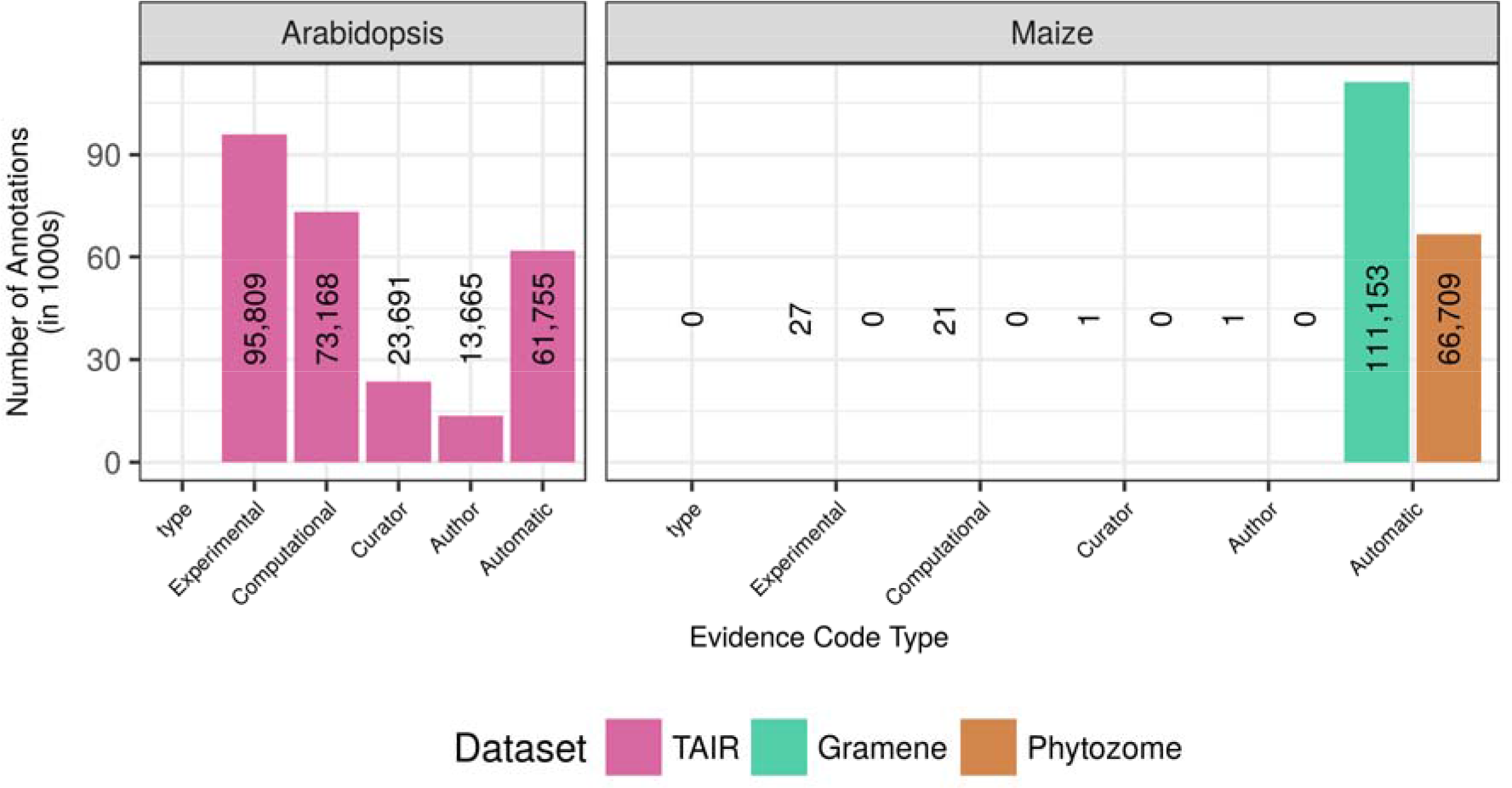
Numbers of annotations by EC category. Arabidopsis (TAIR10) shown in magenta, maize in green and orange for Gramene and Phytozome, respectively. Annotation counts on the y-axis are shown in thousands. Each bar in the histogram is labeled with the actual count to show where counts are so small that no bar is visible.

Exacerbating this problem, transfers of predicted function often are based on sequence similarity alone with no restriction of input data to associations based on well-documented EC types. Furthermore, although mixed-method pipelines like the Ensemble COMPARA pipeline used by Gramene and the Phytozome Pfam2GO (Goodstein et al., 2012; Herrero et al., 2016) mappings may seem reproducible in principle given that they are based on the use of specific systems and software, details including input files and parameters often are unavailable or incomplete, making it impossible for research groups outside the group that generated those annotation resources to reproduce the annotation sets. In addition, because many computational pipelines inherit functional annotations that were also purely computationally derived, a single errant annotation can be propagated to many genomes (Andorf et al., 2007), making it mistakenly appear that many genomes agree on the errant function. For these reasons, existing computational functional annotations of maize (and many other plant genomes) should be approached with skepticism.

Given these issues with the maize functional annotation, we endeavored to create an improved annotation set. This task requires both application of robust and reproducible methods and a gold standard set of maize GO annotations to compare generated result sets to each other as well as to the Gramene and Phytozome maize functional annotations. One small dataset of well-curated GO-based functional annotations does exist for maize. It was initially created by curators at MaizeGDB for the purpose of enriching the MaizeCyc metabolic pathway database (Monaco et al., 2013) and expanded through manual literature curation. This dataset constitutes 1,621 genes and 2,002 GO terms.

We annotated the maize B73 RefGen_v3 annotation set 5b+ using only experimentally-based annotations by filtering out GO assignments with IEA, NAS, and ND ECs from the input data and assigned GO terms using multiple input datasets then compared the performance of sequence-similarity, domain-presence, and mixed-methods based on how well the methods predicted function for genes included in the MaizeGDB gold standard dataset. For mixed-methods, we used pipelines developed for the Critical Assessment of Functional Annotation (CAFA) challenge, a competition designed to evaluate the latest computational functional annotation methods and to promote improvement of methods for functional annotation (Jiang et al., 2016; Radivojac et al., 2013). Groups competing in the CAFA challenge create tools that are are applied to a set of specified target sequences. GO assignments are subsequently evaluated based on accumulation of functional data in the literature for the target sequence set. Some CAFA tools use pre-processing steps combined with a number of different computational and statistical approaches to reduce the number of false positive and false negative annotations (Clark and Radivojac, 2011; Falda et al., 2012; Koskinen et al., 2015). Some mixed-method pipelines performed better on average than other methods in the first iteration of the CAFA competition (Radivojac et al., 2013), indicating that the use of mixed-method pipelines for large scale GO annotations could potentially improve the overall quality of the annotation sets.

The project to evaluate and improve maize GO annotations is called GAMER: <u>G</u>O <u>A</u>nnotation <u>M</u>ethod, <u>E</u>valuation, and <u>R</u>eview. We compared GAMER annotations to annotations based on sequence-similarity, domain, and three CAFA mixed-methods. Next we combined GAMER outputs to generate an aggregate maize-GAMER GO annotation set and compared it to the existing Phytozome and Gramene GO annotations based on the *hF*_1_ score. The GAMER annotations had three major advantages compared to the Gramene and Phytozome annotations: (1) an increased number of maize genes annotated with GO terms; (2) more than twice the number of annotations (GO terms assigned) for maize protein coding genes; (3) similar or better quality scores relative to existing annotations sets based on *hF*_1_ score. The B73 RefGen_v3 5b+ maize-GAMER functional annotation dataset described here is accessible via MaizeGDB (http://download.maizegdb.org/maize-GAMER) and CyVerse (doi.org/10.7946/P2S62P. Scripts used to generate the annotation are available via GitHub at https://github.com/Dill-PICL/maize-GAMER.

*(Note for reviewers: the annotations will also be made available via each MaizeGDB gene model page upon publication.)*

## 3 Materials and Methods

### 3.1 Functional Annotation of Maize Genes

Three sequence-based approaches were used to annotate function to genes in the maize reference genome: sequence-similarity, domain-based, and mixed-method pipelines (see Figure 2; also described in sections 2.1.1, 2.1.2 and 2.1.3, respectively). The scripts (bash, R and Python) used to generate the annotations for maize B73 RefGen_v3 are available at https://github.com/Dill-PICL/maize-GAMER. These scripts run free and open-source tools on different inputs required for these tools to generate annotation datasets. Please refer to the reproducibility supplemental file for details on versions of software, version of input datasets used and commands and parameters used to run these tools.

**Figure 2:**
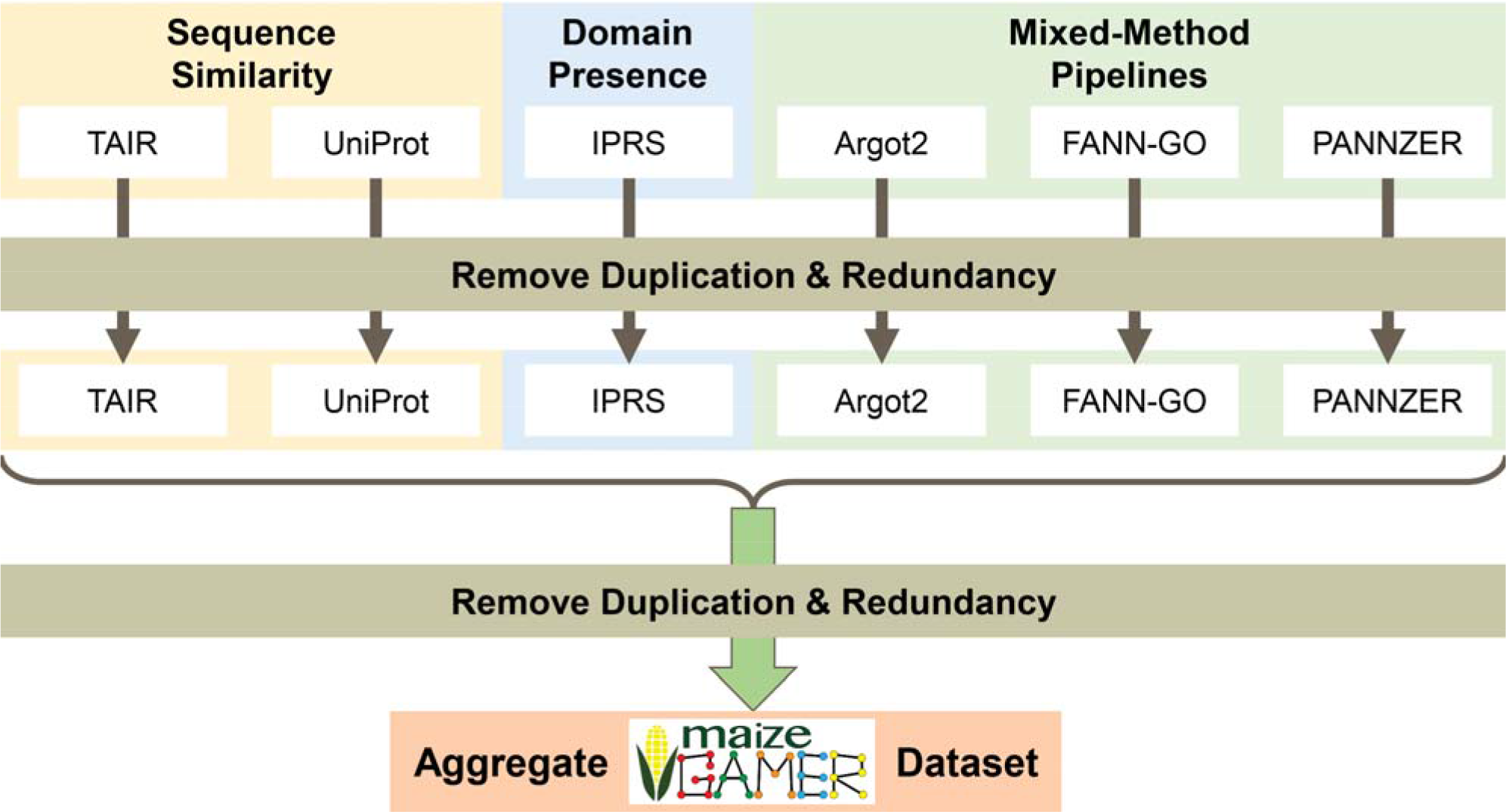
Overview of steps to produce the maize-GAMER datasets. Three types of methods are used: sequence-similarity (yellow), domain presence, (blue) and CAFA mixed-method pipelines (green). Within sequence-similarity, two input datasets were subjected to reciprocal-best-hit against maize: TAIR10 (Arabidopsis) and UniProt (the ten most well-annotated plant species). For domain presence, InterPro signatures were applied to maize using InterProScan (IPRS). From the CAFA mixed-method pipelines, Argot2, FANN-GO, and PANNZER were applied to maize. For each individual output, duplications and redundancies were removed, then the datasets were combined. A second round of duplication and redundancy removal was carried out to produce the maize-GAMER Aggregate dataset.

The B73 genome and protein sequences for gene models included in the Filtered Gene Set (FGS) were downloaded from Gramene Release 42 (Tello-Ruiz et al., 2016). The downloaded protein FASTA file contained sequences for all FGS transcripts (e.g., the gene model X has transcript models X_T01, X_T02, and X_T03). For each gene model only the longest translated protein sequence derived from the transcripts was analyzed. The gold standard annotations used for evaluations were obtained from MaizeGDB, and they encompass GO annotations for 1,619 gene models from RefGen_v3. The number of annotations for cellular component (CC), molecular function (MF), and biological process (BP) were 1,584, 88, 323 respectively.

### 3.2 Sequence-similarity based annotation

The sequence-similarity based annotation method has three main steps: 1) calculation of sequence-similarity, 2) valid hit detection, and 3) inheritance of high-confidence GO annotations. BLASTP was used (Altschul et al., 1990) with default parameters to calculate sequence-similarity between maize protein sequences and two other datasets: the “Arabidopsis” dataset from TAIR, The Arabidopsis Information Resource (Berardini et al., 2015) and the “Plant” dataset from UniProt (UniProt Consortium, 2015). Valid hits were detected using the RBH method from BLASTP results. GO terms with non-reviewed ECs (i.e., IEA, NAS, and ND described in the introduction) were removed from input datasets. All others were inherited between the RBH pairs of maize and the other plant.

Arabidopsis has the largest number of reviewed (human curated) EC GO annotations among plant model organisms (see Supplementary Table S1). A FASTA file of Arabidopsis protein sequences along with the cognate GO Annotation File (GAF) were downloaded from TAIR v.10 (Berardini et al., 2015). The TAIR protein file contained predicted protein sequences from all transcripts. This file was filtered to retain only the protein sequence derived from longest transcript for each gene. Retained protein sequences from TAIR were used to create the TAIR BLAST database, and maize protein sequences were used to create a maize BLAST database. Maize protein sequences were used to query the TAIR BLAST database. Likewise, TAIR sequences were used to query the maize BLAST database. Results from both searches were used to detect RBH pairs between Arabidopsis and maize. All non-reviewed EC GO annotations were removed, and remaining GO associations to Arabidopsis genes were inherited to maize genes for each RBH pair. This maize/Arabidopsis RBH ortholog dataset is called “maize-TAIR GO annotations”.

All reviewed EC GO annotations and protein sequences for all plants from the UniProt-GOA database were downloaded using the QuickGO tool hosted at EBI (Binns et al., 2009). Protein sequences and reviewed EC GO annotations were downloaded separately. The UniProt plant GO annotation dataset containing 304,426 annotations from 75,537 unique protein sequences. The protein sequences downloaded spanned 292 taxa. Only ten species had more than 1,000 annotations (see Supplementary Table S1). Annotations from the top 10 species (in terms of number of reviewed GO annotations) were retained for our analyses. The process to annotate maize genes using UniProt plant data was similar to that for Arabidopsis. Maize protein sequences were matched against protein sequences from each species separately using BLASTP. Putative orthologs were determined using RBH for each maize-plant pair. Terms annotated to the other plant protein were inherited to the maize protein sequence for each putative ortholog pair. GO annotations inherited from each plant species were concatenated together. The derived dataset is called the “maize-UniProt GO annotations”.

### 3.3 Domain Presence

InterProScan5 (IPRS) version 5.16-55.0 was used to create domain based GO annotation of maize protein coding genes (Jones et al., 2014). IPRS was used to annotate GO terms to maize genes to produce the “maize-IPRS GO annotations”.

### 3.4 Mixed-Method Pipelines

At the beginning of this project, the first iteration of the CAFA challenge (CAFA1; described in the Introduction) had been completed. The results from the challenge indicated that CAFA1 mixed-method pipelines performed as well or better than standard methods (Radivojac et al., 2013). To determine their predictive power for functional annotation in plants, the top-performing mixed-method pipelines from CAFA1 were reviewed to identify a group that could be implemented based upon availability of code and sufficient documentation. Three tools were selected: Argot2, FANN-GO, and PANNZER (Clark and Radivojac, 2011; Falda et al., 2012; Koskinen et al., 2015).

#### 3.4.1 Argot2

Argot2 has a batch processing tool that can annotate up to 5,000 pre-processed input sequences. There are two different pre-processing steps for Argot2: 1) querying the UniProt database for sequence-similarity matches to the input sequences, and 2) querying the the Pfam database for putative domains present in the input sequences. The maize sequences were split into multiple FASTA files containing a maximum of 5,000 sequences. The eight FASTA files resulting from the previous step were used to query the UniProt database using BLASTP for matches and the output was saved. HMMER was used to search a local Pfam database for potential hits for all the input protein sequences (Finn et al., 2011). Preprocessing each input FASTA resulted in a pair of input files for Argot2: BLAST and HMMER files. Each pair of pre-processed files was compressed and submitted to Argot2 batch processing tool. Results from each pair of pre-processed data were downloaded and concatenated to create the “maize-Argot2 GO annotations”.

#### 3.4.2 FANN-GO

The file containing maize protein sequences was imported into MATLAB using a built-in function (MATLAB:2017). The MAIN function from FANN-GO was used to pre-process and annotate maize protein sequences. FANN-GO uses BLASTP to query FANN-GO training sequence dataset (derived from UniProt) for potential matches for the input sequences and converts the results to input feature vectors. The FANN-GO predictor built from the training dataset is then used to process the input feature vectors and calculate the probability that a particular protein is associated to a particular GO term. These probabilities are represented in a matrix where rows represent sequences and columns represent GO terms. The matrix was converted to a GAF (GO Annotation File Format) file to be used for subsequent evaluations. This dataset is referred to as the “maize-FANN-GO annotations”.

#### 3.4.3 PANNZER

Maize protein sequences were pre-processed using BLASTP to query a local UniProt protein BLAST database, and the output was saved in XML format as required by PANNZER. PANNZER was run on the xml file output from the previous step, and the output was converted into a GAF file. This dataset from PANNZER is referred to as the “maize-PANNZER GO annotations”.

### 3.5 Metrics Used in maize-GAMER

A number of metrics defined and described by the AIGO (Analysis and the Inter-comparison of GO functional annotations) library were used to select high-confidence annotations, clean, and evaluate the maize-GAMER derived annotation sets (Defoin-Platel et al., 2011). AIGO has defined two type of metrics: analysis metrics and comparison metrics (see Table 2).

**Table 2:**
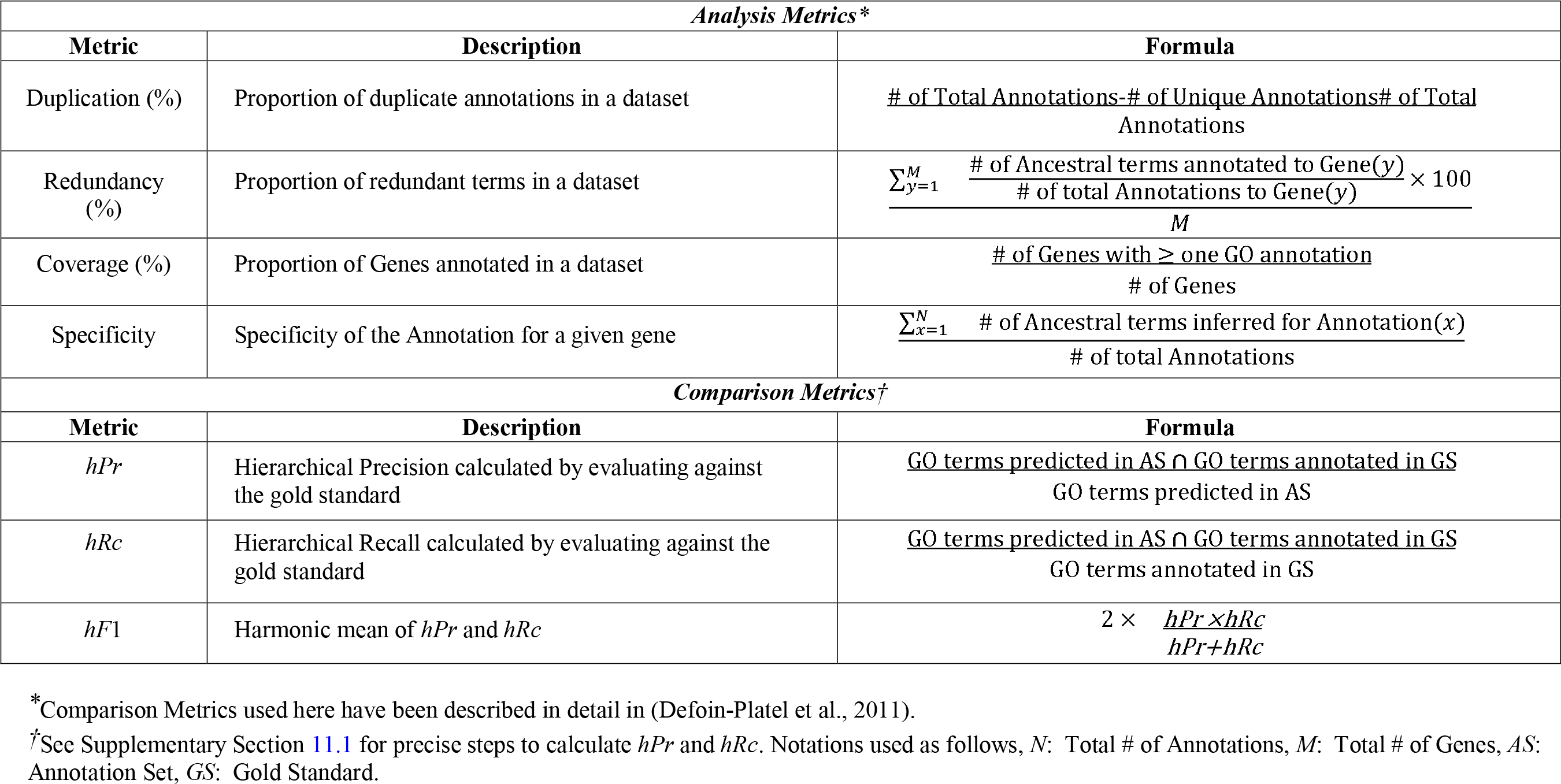
Metrics used to analyze and evaluate the annotation datasets

#### 3.5.1 Analysis Metrics

Analysis metrics defined by AIGO measure features of a given annotation set. Four AIGO analysis metrics were used for maize-GAMER: Duplication, Redundancy, Coverage, and Specificity (see Table 2). With respect to calculating AIGO metrics, an annotation is defined as a single gene-GO term pair. Duplication is the proportion of annotations that are not unique to a given annotation set. Duplication is calculated for each annotation set as described in Table 2. The collection of more general GO terms that can be inherited from a specific GO term are called ‘ancestral’. Redundancy occurs when a GO term and one or more of its ancestral terms are annotated to the same gene in a dataset. Redundancy metric was calculated by obtaining the mean of the proportion of ancestral terms annotated for each gene in a particular dataset. Coverage is the proportion of genes that have at least one GO term assigned. Specificity is measured by counting the number of ancestral terms that exist for a given annotation, then averaging those counts across all annotations in the set. An ideally cleaned annotation dataset would have no duplication, no redundancy, high coverage, and high specificity.

#### 3.5.2 Comparison Metrics

Comparison metrics defined by AIGO measure how well a given set of annotations match with another set of annotations. The AIGO comparison metrics hierarchical Precision (*hPr*) and hierarchical Recall (*hRc*) were used to evaluate annotation sets against gold standard annotations from MaizeGDB (see Table 2 & Supplementary Materials). Different metrics have been defined for the evaluation of GO annotations against a gold standard (Clark and Radivojac, 2013; Defoin-Platel et al., 2011; Jiang et al., 2016; Radivojac et al., 2013). AIGO provided a set of well described evaluation metrics which were adapted by maize-GAMER and was utilized for large number of annotations produced by mixed-method pipelines. Both *hPr* and *hRc* evaluations start with propagating the GO terms in the annotations to the root.

*hPr* is the proportion of the GO terms (directly annotated and inferred by propagation) in an annotation set which is shared with the GO terms (directly annotated and inferred by propagation) in the gold standard. *hRc* is the proportion of the GO terms in the gold standard which are found in the annotation set. *hPr* and *hRc* were calculated for the genes in the gold standard dataset, and were calculated independently for each GO category. If a gene was annotated in the gold standard, but was not annotated in the annotations set then both *hPr* and *hRc* were set to 0. See supplementary materials for precise steps used to calculate *hPr* and *hRc.* Harmonic mean (*hF*_1_) of *hPr* and *hRc* was calculated for each annotation set for each GO category to use a single number to compare different annotation methods.

### 3.6 Cleaning and Combining Component Datasets

#### 3.6.1 Score Threshold Selection for Mixed-Methods

Mixed-method pipelines used in the maize-GAMER project provide a confidence score for each GO annotation. The confidence score ranges from 0.0-1.0, where a higher score indicates more confidence for a given annotation. A score threshold which maximizes *hF*_1_ (*hF_max_*) will select the optimal set of annotations which reduces the total number of false-positives and false-negatives (see section 2.2 for description of the metrics). The range of annotation scores from mixed-method pipelines did not span the whole 0.0-1.0 range, so the scores were normalized to fall between 0.0-1.0 independently for each annotation set. A set of thresholds (every 0.05 from 0.0 to 1.0; i.e. 0.00, 0.05, 0.1, 0.1, …., 0.95, 1.00) were selected. *hF*_1_ score for each GO category and each threshold was calculated by selecting the annotations with a normalized score that was ≥ to the threshold and evaluating against the gold standard annotations. *hF_max_* for each GO category was determined by getting the highest *hF_1_* obtained from the previous step. The score thresholds which resulted in *hF_max_* were used to select a subset of annotations from each mixed-method pipeline (see Supplementary Table S2). The maize-Argot2, maize-FANN-GO, maize-PANNZER GO annotations described in subsequent sections refer to the subset of annotations selected via this selection step.

#### 3.6.2 Removing Redundancy and Duplication

Duplication is the presence of two or more instances of the same gene-GO term pair in a single annotation set. Redundancy is the presence of an ancestral GO term in the annotations of a gene which also contains a specific annotation from which the ancestral GO term can be inferred by propagation. Component annotation sets from all methods described above were cleaned by removing redundancy and duplication for each annotation set across all three GO categories. Duplication was cleaned by replacing multiple instances of a gene-GO term pair with a single instance for a given annotation set. Duplicate annotations from all six raw annotation sets were removed and files with non-duplicate annotations were created for each annotation set. Redundancy was cleaned by removing annotations containing GO terms that could be inferred from other terms based on the GO hierarchy, and only retaining the annotations with GO terms that cannot be inferred.

#### 3.6.3 The maize-GAMER Aggregate Dataset

Clean (non-redundant and non-duplicated) annotation sets from all component methods were merged to generate the maize-GAMER aggregate annotation set. Redundancy and duplication introduced by concatenating multiple datasets were removed.

A new genome assembly (B73 RefGen_v4) and annotation set (Zm00001.2) for maize inbred line B73 was recently released (Jiao et al., 2017). Because this dataset has not been available for long, only few published analyses are available and the research community is only now in the process of transitioning to general use of RefGen_v4 for large-scale analyses. As such, analyses and results described here derive from the well-annotated v3 assembly and annotation set. To extend outcomes of the work described here for future v4 efforts, maize-GAMER aggregate annotations have also been created for the maize B73 RefGen_v4, which can be accessed at MaizeGDB (http://download.maizegdb.org/maize-GAMER) and via CyVerse (doi.org/10.7946/P2M925).

### 3.7 Evaluation of GAMER-derived Annotation Sets

Component and aggregate annotation sets were compared at two levels; a general comparison, and a GO category-specific comparison.

Comparison metrics mentioned in Section 2.2 were calculated for the general comparison (see Table 2). All metrics were calculated independently for each annotation set, and compared among component annotation sets as well as the aggregate annotation set. Coverage and the number of annotations were calculated directly for each annotation set. Specificity was calculated for each annotation and the mean across all annotations is reported.

The annotations from component annotation sets and the aggregate annotation set were divided into specific GO categories and category-specific annotations were evaluated separately. Three different metrics were used for GO category-specific evaluations: coverage, number of annotations, and *hF*_1_ (see section 2.2 for more details). Coverage and the number of annotations were calculated individually for each GO category for each dataset. *hF*_1_ score was calculated for each annotation set for each GO category.

### 3.8 Comparisons among the maize-GAMER Aggregate, Gramene, and Phytozome Annotation Sets

The existing Gramene, Phytozome, and maize-GAMER annotations were compared to each other. Redundancy and duplication were removed from the Gramene and Phytozome annotation sets before evaluations were performed. Evaluation and comparisons were identical to the analyses performed in the previous section 2.4. General evaluations for the maize-GAMER, Gramene, and Phytozome annotation sets were based on coverage, number of annotations, and specificity. These metrics were calculated as described in the previous section 2.4. The Gramene, Phytozome, and maize-GAMER annotation sets were also compared in a GO category-specific manner to account for biases in performance among different categories (i.e., CC, BP, and MF). Comparisons were made based on coverage, number of annotations, and mean *hF*_1_ score.

### 3.9 Case Study of the Gene *nana plant1 (na1)*

The gene *na1* (GRMZM2G449033) had the most terms (7 GO terms) associated with it in the gold standard dataset, and all the terms were from BP GO category. Annotations for *na1* from the three maize annotation sets were obtained. The ancestral nodes were inferred from the leaf nodes for each annotation set, and a subgraph for the BP ontology was generated (see Fig 5). The nodes in the subgraph were compared to gold standard and nodes shared between a given annotation set and the gold standard dataset were identified. Nodes exclusively found only in a given annotation set or the gold standard were also identified. Illustrations of the subgraphs without node labels were drawn to compare among the three different GO annotation sets.

**Figure 5:**
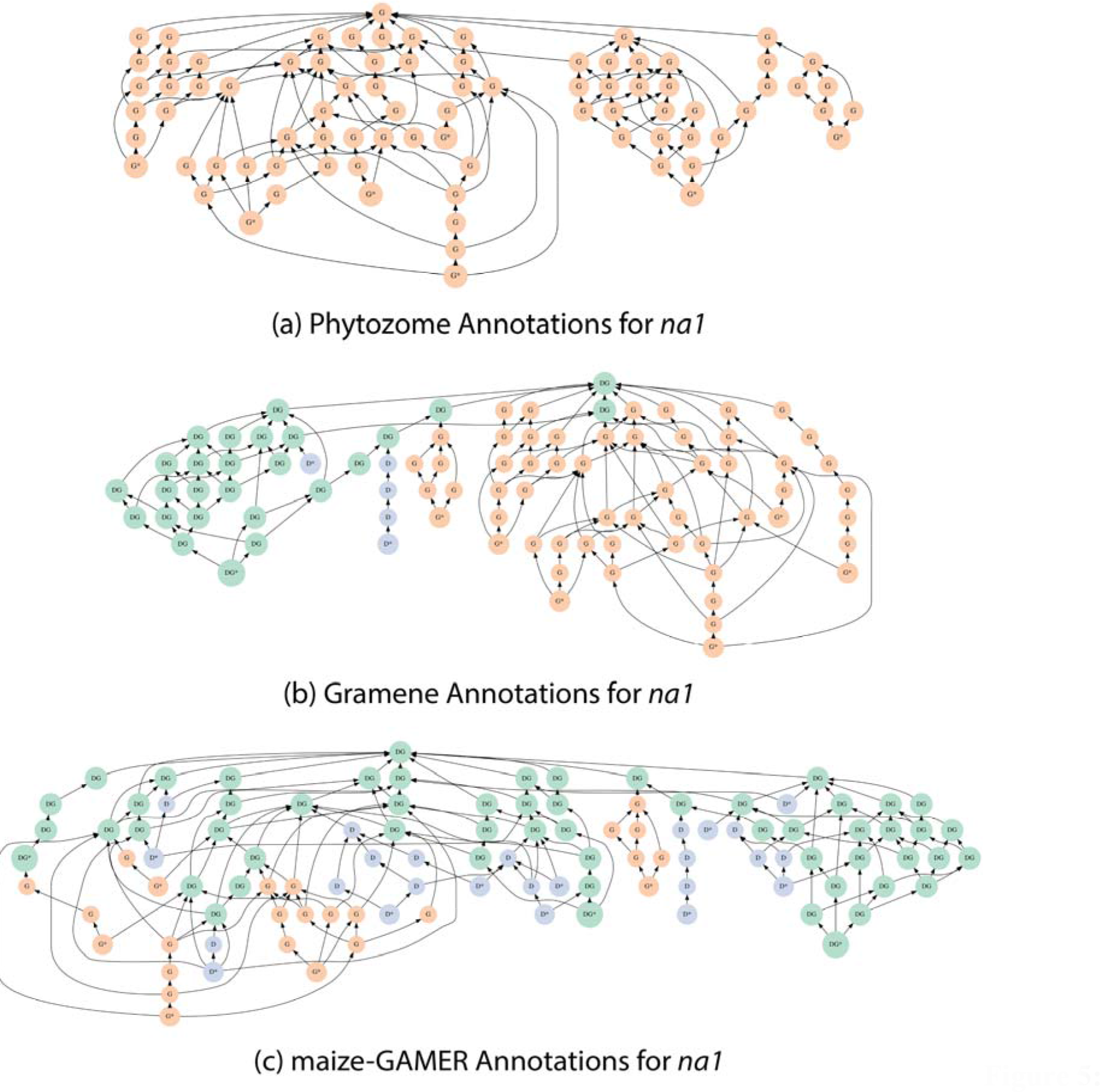
Biological Process GO graph for maize *na1*. Leaf terms are toward the bottom, root terms are toward the top. Terms covered only by the gold standard are shown in orange (labeled G), those in the dataset but absent from the gold standard are shown in blue (labeled D), and those that appear in both are shown in green (labeled DG). Leaf terms in each subgraph have an * next to them. Phytozome graph is shown at the top (5a) Gramene graph is shown in the middle (5b), and maize-GAMER aggregate graph is shown at the bottom (5c).

## 4 Results

### 4.1 Evaluation of maize-GAMER Derived Component Annotation Sets

The maize-GAMER derived component annotation sets (i.e., the TAIR, UniProt, IPRS, Argot2, FANN-GO, and PANNZER) and the maize-GAMER aggregate annotation set were evaluated across GO categories as well as within each GO category using metrics described in Table 2.

#### 4.1.1 General Evaluation of maize-GAMER Component Annotation Sets

Initial evaluations and comparisons of datasets created by the maize-GAMER pipeline were assessed based on coverage, number of annotations, and specificity among all clean component annotation sets as well as the aggregate annotation set (see Tables 2 & 3). The TAIR and UniProt annotation sets had the lowest coverage and number of annotations among all maize-GAMER component annotation sets (Table 3). The Argot2, FANN-GO, and PANNZER annotation sets had the highest number of annotations compared to other annotation sets, as well as higher coverage compared to other annotation sets. Notably, FANN-GO had the highest coverage at 100% of genes, and Argot2 had annotations for more than 90% of the genes. The IPRS annotation set had a lower number of annotations compared to the CAFA mixed-method pipelines, but covered more genes than sequence-similarity methods. Although sequence-similarity methods and IPRS covered a lower number of genes, they had higher specificity compared to mixed-method pipelines in general. Of the three mixed-method pipelines, only PANNZER had comparable specificity to the methods, but had lower coverage than both Argot2 and FANN-GO. Both Argot2 and FANN-GO had lower average specificity, but had higher coverage than other methods. The maize-GAMER aggregate annotation (made up of all component annotation sets) covered all maize genes with at least one annotation (as expected given that the FANN-GO component annotation set also covers 100% of genes). In addition, the aggregate annotation set contains more than double the number of annotations that occur in any component annotation set. This indicates that different component methods assign different GO terms to genes. Therefore, combining annotations from different methods results in increased diversity of GO term assignments. Moreover, the aggregate annotation set has higher specificity than the mixed-method pipelines which have higher coverage, but has lower specificity than all other component annotation sets.

**Table 3:** Results from the general evaluation of maize-GAMER derived annotation sets

| Method Type | Sequence-Similarity |  | Domain | CAFA Mixed-Methods |  |  | Union |
| --- | --- | --- | --- | --- | --- | --- | --- |
| Annotation Set | TAIR-RBH | Plant-RBH | IPRS | Argot2 | FANN-GO | PANNZER | Aggregate |
| Raw Annotations | 61,528 | 106,053 | 200,324 | 450,013 | 618,312 | 1,091,123 |  |
| Duplication (%) | 9.11 | 65.89 | 72.18 | 0.33 | 1.07 | 0.97 |  |
| Redundancy (%) | 14.18 | 9.17 | 10.91 | 48.25 | 59.84 | 73.09 |  |
| Clean Annotations | 35,791 | 32,085 | 46,599 | 224,827 | 187,850 | 219,984 | 515,059 |
| Coverage (%) | 23.70 | 20.10 | 48.48 | 92.64 | 100.00 | 47.86 | 100.00 |
| Specificity | 11.54 | 12.20 | 10.45 | 8.54 | 5.30 | 11.58 | 9.56 |

Genes that are annotated with at least one GO term from each component annotation set were compared among the three different method types (i.e., sequence-similarity, domain-based, and mixed-methods; see Fig 3a). This comparison revealed that less than a quarter of genes had been annotated by all three methods, but more than half were annotated by two different methods. The remainder were only annotated by mixed-method pipelines. Sequence-similarity and domain-based methods resulted in annotations to genes that were also annotated by mixed-method pipelines. The number of genes annotated by domain-based methods and mixed-method pipelines, and are not annotated by sequence-similarity based methods are higher than genes annotated by all three methods. In contrast, sequence-similarity methods shared more genes with both other annotation sets than only with mixed-method pipelines. Moreover, only mixed-method pipelines annotate at least one GO term to all genes in the maize FGS.

**Figure 3:**
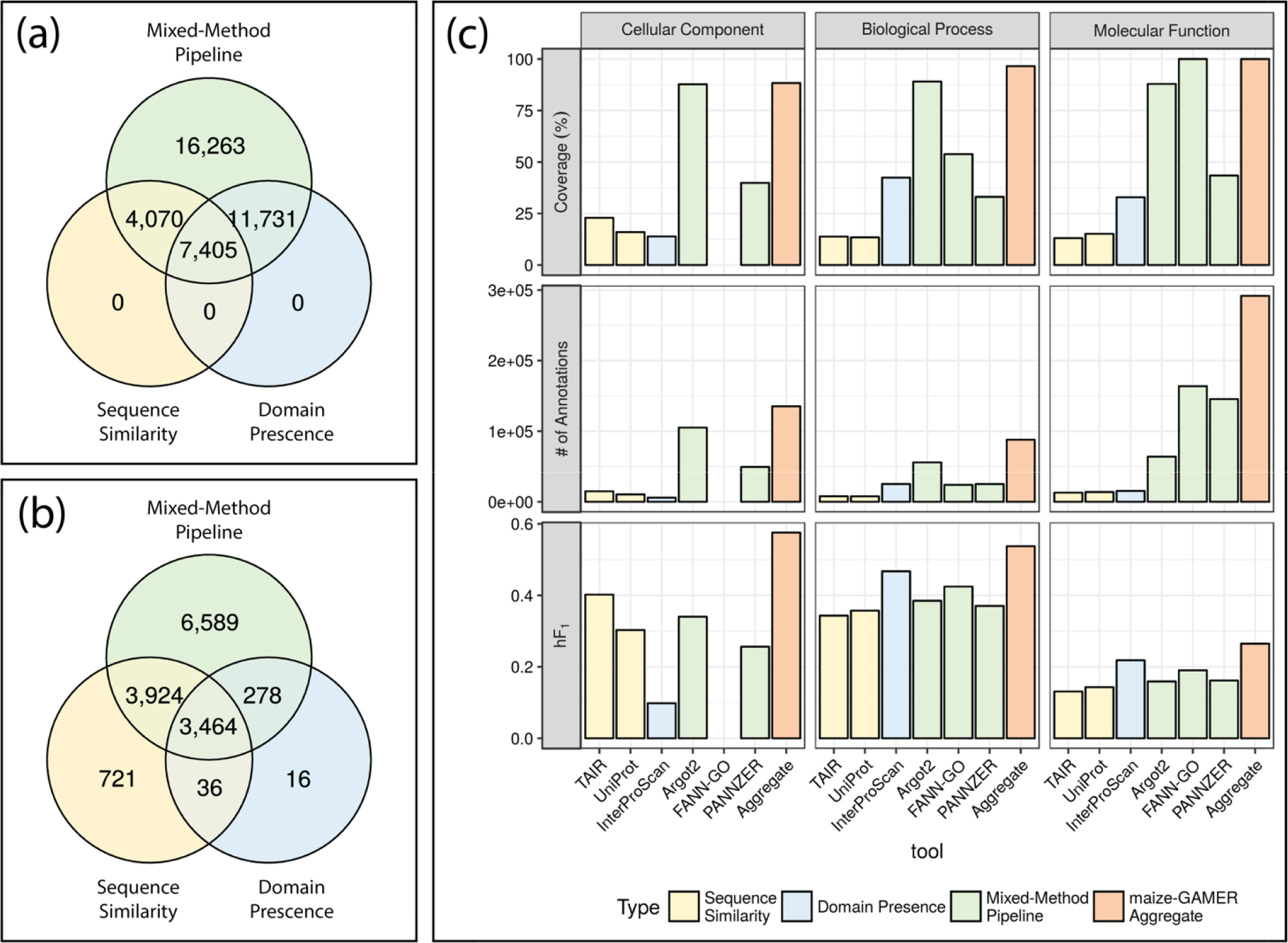
GO assignment metrics for each method type. Sequence-similarity in yellow, domain presence in blue, and mixed-method pipeline in green. (a) Number of genes with at least one GO term annotated. (b) Number of GO terms with at least on gene annotated. (c) Percent coverage, number of annotations, and average *hF*_1_ score for each annotation set across the three GO graphs (i.e., Cellular Component, Molecular Function, and Biological Process). Color codes as used in (a) and (b), with the aggregate dataset shown in orange.

Although the mixed-method pipelines annotated all genes, they did not capture all GO terms annotated to genes by the other methods. GO term assignments were compared to evaluate the diversity of the GO terms present in the three types of GO annotation methods (see Fig 3b). GO terms annotated directly and ancestral terms inferred from the direct terms annotated to genes were compared among the three GO annotation methods used in maize-GAMER. The number of GO terms annotated by sequence-similarity, domain-based, and mixed-method pipelines were 3,794, 8,145, and 14,225, respectively. The number of GO terms annotated by the mixed-method pipelines are significantly higher than both other methods, however there are a small number of GO terms that are only annotated by sequence-similarity (721) and domain-presence (16) methods. Only a small proportion (23.05%) of the total (15,028) GO terms are annotated by all three methods.

#### 4.1.2 GO Category-specific Evaluations of maize-GAMER Component Annotation Sets

CAFA1 indicated that annotations for some GO categories are easier to predict than others (Radivojac et al., 2013). This indicated that the GO category specific evaluations could provide a more accurate comparison between component methods. This would also allow unbiased comparison of tools which do not predict certain categories (e.g., FANN-GO doesn’t predict the CC category). Therefore, maize-GAMER derived annotation sets were divided into specific GO categories (i.e., CC, BP, and MF) and each category was evaluated separately based on coverage, number of annotations, and *hF*_1_.

Mixed-method pipelines had higher coverage across all three GO categories. Argot2 covered more than 80% genes across all three categories (see Fig 3c). FANN-GO does not annotate GO terms for CC category, but had 100% coverage in BP category, and covered about 50% genes in MF category. PANNZER had the lowest coverage compared to the other mixed-method pipelines, and covered only 30-50% of genes across different categories, and had highest coverage in BP. Sequence-similarity methods consistently had lowest coverage compared to other methods in BP and MF, but IPRS had the lowest coverage in CC. IPRS covered higher number of genes than sequence-similarity methods in BP and MF, but had lower coverage than mixed-method pipelines. When comparing IPRS coverage across three GO categories, the coverage was highest in MF. Aggregate annotation set covered slightly more genes than the component annotation sets with highest coverage in each category, and covered more than 88% of the FGS genes in all categories. In the BP category, the aggregate annotation set annotated all genes from maize FGS with at least one annotation.

Mixed-method pipelines produce a higher number of annotations than other methods in all three GO categories. Moreover, the number of annotations from mixed-method pipelines loosely correlate with coverage in different GO categories. The only exception was PANNZER, which annotated more GO terms per gene in BP category (data not shown), than any other component annotation set. The number of annotations from sequence-similarity methods and IPRS were consistently lower than mixed-method pipelines. The variation in the number of annotations was proportional to the number of genes annotated in sequence-similarity and IPRS methods. The lowest number of annotations was seen in the CC category from IPRS, and sequence-similarity methods in other GO categories. As the union of all component method annotations, the aggregate annotation set had a higher number of annotations in all three GO categories. The highest number of annotations for the aggregate annotation set was from the BP GO category, followed by CC and MF.

We used *hF*_1_ scores as a representation of the quality of annotations and annotation sets. As described in materials and methods, the *hF*_1_ was calculated individually for all genes in the gold standard dataset and then averaged across all genes within an annotation set. There are clear differences in *hF*_1_ across different GO categories. The highest performance was seen in the MF category, and the lowest performance is seen in the BP category. This fits the observation from CAFA1 (Radivojac et al., 2013). Mixed-method pipelines outperformed other methods in all three GO categories. PANNZER produced the highest *hF*_1_ within the MF category, but Argot2 had the highest *hF*_1_ scores in CC and BP. IPRS outperformed sequence-similarity methods in both MF and BP categories, but was the lowest performing method in the CC category. Comparison between two sequence-similarity methods indicated that maize-UniProt method performs better than the maize-TAIR method in MF and BP categories. On the other hand, maize-TAIR method performs better than maize-UniProt method in the CC category. Aggregating the component annotations from maize-GAMER increased the performance in the CC category. In contrast, aggregating the component annotation sets did not increase the performance compared to the top performing tool in other categories.

### 4.2 Evaluation of Existing Maize GO Annotation Sets and Comparison to the maize-GAMER Aggregate Annotation Set

Two existing maize GO annotation sets, Gramene and Phytozome, were downloaded, evaluated, cleaned (i.e., redundancies and duplicates were removed), and compared with maize-GAMER aggregate annotations (referred to as the “maize-GAMER annotation set”). The same metrics used for the evaluation of maize-GAMER derived annotation sets were used for the comparison among the existing maize GO annotation sets and maize-GAMER aggregate annotation set.

#### 4.2.1 General Evaluation of Public Maize GO Annotation Sets

The maize-GAMER aggregate annotations covered all genes in the maize FGS with at least one GO term, but Gramene and Phytozome covered only about half the genes (see Table 4). Phytozome covered the fewest genes (less than half of the genes), and Gramene covered slightly more than half of the genes (see Table 4). The maize-GAMER annotation set had more annotations than both Gramene and Phytozome. Gramene had two-fold more annotations than Phytozome, and the maize-GAMER had severalfold more annotations than Gramene. While the maize-GAMER has higher coverage and a higher number of annotations, it has lower average specificity than Gramene and Phytozome. Gramene has the highest average specificity of all three annotation sets.

**Table 4:** Overall results from maize-GAMER and other existing maize datasets

| Analysis Metric | Existing |  | maize-GAMER |
| --- | --- | --- | --- |
|  | Gramene | Phytozome | aggregate |
| Raw Annotations | 111,203 | 66,709 |  |
| Duplication (%) | 0.00 | 37.90 |  |
| Redundancy (%) | 23.25 | 7.24 |  |
| Clean Annotations | 81,315 | 36,987 | 515,059 |
| Coverage (%) | 55.55 | 40.87 | 100.00 |
| Specificity | 10.90 | 10.41 | 9.56 |

Genes with annotations from each set were compared to see the distribution of annotated genes among different annotations (see Fig 4a). Genes from Gramene and Phytozome annotations were a subset of the maize-GAMER annotations. Less than half of the genes were annotated in all three sets, and slightly more than half of the genes were annotated in at least two sets. Comparison of Gramene and Phytozome annotations show that most of the genes that were annotated were shared. Both Gramene and Phytozome had genes that were annotated in only one of the two (i.e., Gramene or Phytozome but not both; See Fig 4a).

**Figure 4:**
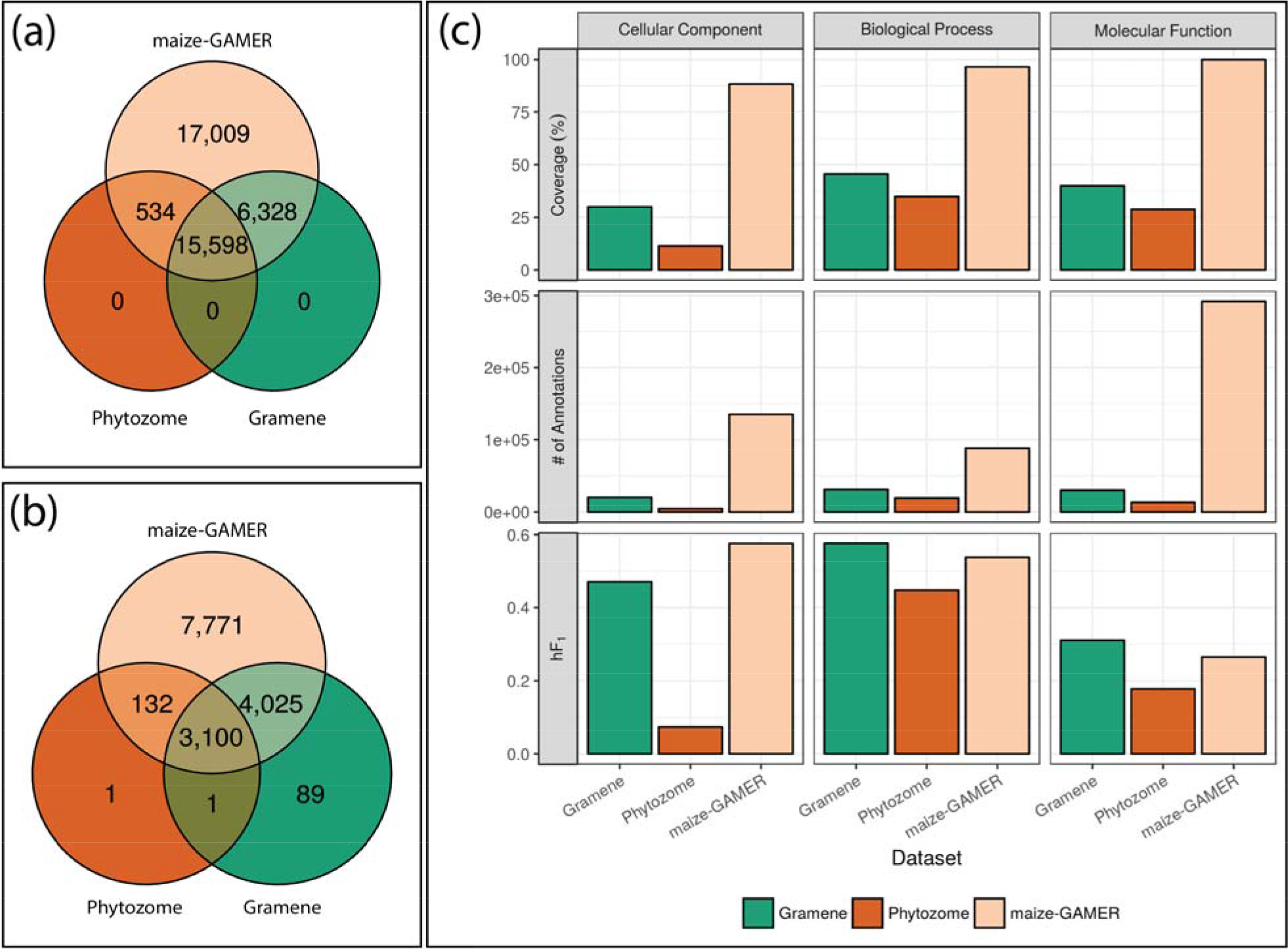
GO assignment metrics for Gramene, Phytozome, and maize-GAMER. Gramene in green, Phytozome in rust, and maize-GAMER in tan. (a) Number of genes with at least one GO term annotated. (b) Number of GO terms with at least one gene annotated. (c) Percent coverage, number of annotations, and *hF*_1_ score for each annotation dataset across the three GO graphs (i.e., Cellular Component, Molecular Function, and Biological Process).

GO terms annotated directly to genes by different methods and ancestral GO terms propagated from these annotations were compared among the three annotation sets. The number of GO terms annotated in each set varied greatly. The least diverse set in terms of number of GO terms annotated was Phytozome, which was annotated with only 3,234 GO terms (approximately 7% of total GO terms). Gramene has annotated 7,215 GO terms (approximately 16%), and was more diverse than Phytozome, but had lower diversity than maize-GAMER. maize-GAMER had the highest diversity and contained 15,028 GO terms (approximately 33%). A small number of GO terms were used by all three annotation sets, and the majority of terms from Phytozome were shared across all three annotation sets. Only a single GO term was exclusive to the Phytozome annotations, and small number of terms were found to be exclusive to Gramene annotations. Approximately 50% of the GO terms from maize-GAMER were unique. The maize-GAMER aggregate annotations shared a higher number of GO terms with Gramene than Phytozome.

#### 4.2.2 GO Category-specific Evaluations of maize-GAMER and Existing Maize GO Annotation Sets

Annotations from the three maize GO annotation sets were analyzed in a GO category-specific manner to identify differences in performance among the different categories (see Figure 4c). As was true for the component annotation sets, three different metrics were used for evaluation and comparison: coverage, number of annotations, and *hF*_1_ score.

Comparison of coverage across GO categories indicated that all annotation sets had lower coverage in CC category, compared to other categories. Both Gramene and Phytozome had lower coverage in BP than MF, but maize-GAMER had higher coverage in BP than MF. Lowest coverage for all annotation sets and categories was seen in the CC category for the Phytozome annotation set, and the highest coverage was seen in the maize-GAMER aggregate annotation set in the BP category. Comparison among the three maize annotation sets indicates that the maize-GAMER annotation set had the highest coverage in all three categories by a large margin. Coverage from maize-GAMER was almost twofold that of Gramene, which had the next highest coverage in all GO categories. Gramene had higher coverage than Phytozome in all three categories.

When the number of annotations were compared across different GO categories the lowest number of annotations for Gramene and Phytozome annotation sets were seen in the CC category. In contrast, maize-GAMER had the lowest number of annotations in the MF category. Moreover, both Gramene and Phytozome both had a higher number of annotations in the MF category whereas maize-GAMER had the highest number of annotations in the BP category. Comparison among the annotations sets illustrated that the maize-GAMER annotation set has the highest number of annotations in all three categories. Phytozome had the fewest annotations in all three GO categories. Number of annotations loosely correlated with coverage in different GO categories for both Gramene and Phytozome. Furthermore, maize-GAMER had the highest number of annotations in the BP category. The number of annotations from the maize-GAMER annotations for the BP was severalfold higher than other annotation sets (approximately 9x that of Gramene and approximately 28x that of Phytozome).

The *hF*_1_ score reflects the overall quality of annotations produced by different pipelines used by the three maize annotation projects. Comparing performance of different pipelines across the three GO categories revealed a similar trend that was seen in the previous section. All pipelines had higher *hF*_1_ scores in the MF category and had lower *hF*_1_ scores in the BP category. The only pipeline that did not fit this trend was Phytozome, which had lowest performance in the CC category. maize-GAMER had a higher *hF*_1_ score than other pipelines in the CC category. maize-GAMER also had higher performance than Phytozome in other categories, but performed slightly lower than Gramene in those categories. Gramene performed better than other pipelines in the MF and BP categories. Phytozome consistently had lower performance than other pipelines across all three GO categories. Phytozome’s performance was especially low in the CC category, which was the lowest *hF*_1_ score seen for any annotation set in any GO category.

To visualize how GO annotation methods perform comparatively, the distribution of metrics *hPr, hRc,* and *hF*_1_ can be calculated across all genes included in the gold standard dataset. In Supplementary Figure S1, lower coverage by sequence-similarity and domain-presence methods is illustrated by the high number of genes with a value of zero. In general, mixed-method pipelines are shown to perform better than other methods we used. They cover more genes, and two of the three mixed-method pipelines have higher *hPr* and *hRc* than all other methods. The FANN-GO distribution is different from both other mixed-methods: in general it has lower performance than other methods. This can be attributed to that fact FANN-GO annotations have lower specificity than other methods, and lower specificity results in lower values for *hPr* and *hRc*. In addition, the maize-GAMER has fewer genes with a value of 0 than both Phytozome and Gramene.

### 4.3 Example Annotations from *nana plant1* (*na1*)

The gene *nana plant1* (*na1*; GRMZM2G449033) has more annotations than any other gene in the gold standard dataset. A classical maize mutant with a dwarf phenotype (Hartwig et al., 2011), the *na1* recessive mutant results from a loss-of-function mutation in the gene that affects the brassenosteroid (BR) biosynthetic pathway where BR is a plant hormone that is required for normal plant growth (Hartwig et al., 2011). In the gold standard dataset, *na1* had 7 biological process GO terms annotated. Annotations for *na1* from different maize annotation sets were compared to the gold standard, and a subgraph for each annotation set and gold standard dataset was plotted (see Fig 5). Phytozome did not annotate any GO terms to *na1* (see Fig 5a), but both Gramene (see Fig 5b) and maize-GAMER (see Fig 5c) have annotated BP GO terms for *na1*. Gramene annotates 3 GO terms *na1* while maize-GAMER has annotated 13 GO terms to *na1*. Two GO terms from the gold standard are known to be related to *na1* dwarf phenotype from previous studies, “brassinosteroid biosynthetic process” (GO:0016132) and “unidimensional cell growth” (GO:0009826). While both of these were annotated correctly by maize-GAMER (see Fig 5c), only one of them was correctly annotated by Gramene (see Fig 5b). Comparison of overlapping nodes indicates that the maize-GAMER aggregate annotation set also contains a number of less specific non-leaf terms which overlap with nodes inferred from gold standard dataset. Overall, the maize-GAMER has larger proportion of overlapping nodes with the gold standard than the Gramene for the BP GO category.

The different approaches taken by the pipelines from Gramene and maize-GAMER result in different annotations for the example case study of the maize *na1* gene. Gramene has a lower number of GO terms annotated to *na1* than maize-GAMER. The average specificity of GO terms annotated in the BP category for *na1* (see Figure 5) is not significantly different between GAMER (mean=12.154) and Gramene (mean=12.667) pipelines (2-sided 2-group Wilcoxon rank-sum test; p = 0.89). This example from *na1* indicates that the specificity of the annotations are not significantly different for specific instances, but are different when compared overall. We further compared the GAMER component and aggregate dataset creation methodologies annotate gene function to that of Gramene and Phytozome by comparing the hPr, hRc, and hF_1_ metrics for the *na1* gene (See Supplementary Figure S2). Phytozome’s method stands out because it has no annotations for *na1*, thus all the metrics have a value of 0. The GAMER aggregate dataset has higher *hF_1_* score compared to Gramene, and has higher *hPr* and *hRc* as well. Among the component datasets, Pannzer has the highest *hF_1_* score and highest *hRc*. InterProScan has the highest *hPr,* but its lower *hRc* reduces the hF_1_. FANN-GO and Argot2 have lower metrics due to lower specificity of annotations compared to other methods.

## 5 Discussion

In keeping with our goal, through the maize-GAMER project we were able to improve the GO annotation dataset for maize and to document inputs, methods, and results at a level that enables both reproducibility and reuse of the pipeline for future genome versions.

To determine how best to create an improved maize annotation dataset, we tried out multiple different methods and compared the resulting datasets based on a gold standard set of gene functions. This also enabled us to better understand differences in term assignments among the methods we used. Using the same gold standard, we also were able to compare resulting datasets to those produced by and available from Gramene and Phytozome. We used the *hF_max_* metric to select high-confidence annotations from mixed-method pipelines and to evaluate annotation sets resulting from all methods under evaluation. We found that mixed-method pipelines developed for the CAFA1 challenge outperformed RBH and domain-presence methods for GO annotation (Radivojac et al., 2013). They covered more genes with annotations, produced higher number of annotations, and had higher *hF*_1_ score than both sequence similarity and domain-based methods. The higher performance from mixed-method pipelines are the outcome of advanced statistical (Falda et al., 2012; Koskinen et al., 2015) and machine learning approaches (Clark and Radivojac, 2011) used to reduce the false positive and false negative annotations. Mixed-method pipelines do have a limitation: they have higher coverage but annotations are less specific in general when compared with datasets produced using other approaches. This could be due to the dearth of training dataset for the more specific GO terms, which is required for training machine learning methods.

When we aggregated the predictions from RBH, domain-based methods, and three tools from CAFA1, we produced the maize-GAMER aggregate dataset, which covers more gene space than the datasets produced by Gramene and Phytozome, and with similar or better accuracy. While the higher coverage could cause concern, evaluating the annotations using the gold standard has shown that the performance is similar or better than existing datasets. This also indicates that maize-GAMER annotations are no less reliable than Gramene and are in fact are better than those for Phytozome. Removing less specific GO terms annotated by some methods in cases where there were more specific GO terms annotated to the same genes was important for aggregating different datasets. In certain cases, more than one annotations with lower specificity were replaced by a single annotation with higher specificity. As with any computational approach to annotate GO terms, the current maize-GAMER dataset should be considered as an initial step in improving the GO annotations in maize. As future iterations of the CAFA competition evaluate new tools and methods for GO annotations, we anticipate that the quality of computational maize GO annotations could be iteratively improved in a reproducible manner by continuing to apply the newest, best performing methods.

To enable better reproducibility, we have generated a supplementary document with exact parameters and commands used to generate the maize dataset. We are currently in the process of formalizing the code used to generate the maize GO annotation set into a reusable pipeline called GO-MAP. Once completed, the GO-MAP pipeline can be used for GO annotation of newly sequenced plant genomes as well as existing plant genomes. The pipeline will be made freely available, and will utilize the same methods and datasets used for maize. The set of manually reviewed gene function annotations for maize that we call the gold standard is both incomplete and sparse. This situation does not reflect the amount of published literature describing gene function for maize. Instead, this situation is due to limited curation of gene function into GO terms. While tools exist at MaizeGDB that enable researchers to assign GO terms to genes directly, these tools remain poorly utilized. In an effort to improve community engagement and to upgrade the evidence codes for GO assignments, our next step for maize-GAMER will be to develop and deploy a tool to enable experts in the maize community to review existing GO annotations. By enabling GO annotation review through expert crowdsourcing, term assignments produced by computational pipelines including GAMER can be upgraded from IEA (inferred from electronic annotation) to RCA (reviewed computational analysis). In this way, we will enable the transfer of collective knowledge members of the maize community have generated over the years to produce higher-quality functional annotation datasets for maize with clear extension of this practice for other species.

## 7 Acknowledgements

We thank Darwin Campbell and Ramona Walls for efforts to create the DOIs for datasets available via CyVerse and John Portwood and Ethalinda Cannon for their efforts to populate resulting datasets for MaizeGDB. Thanks to members of the Dill Plant Informatics and Computation Lab (dill-picl.org) for critical review and helpful suggestions, with particular thanks to Darwin Campbell for help with figures.

This work was supported by funding from the Iowa State University Plant Sciences Institute Faculty Scholars Program to C.J.L.D.; the National Science Foundation [IOS #1027527] to C.J.L.D.; and the United States Department of Agriculture - Agricultural Research Services to C.M.A. IF was supported, in part, by National Science Foundation award ABI-1458359.

## 8 Authors’ Contributions

K.W. carried out all of the research and development efforts. K.W., C.M.A., I.F., and C.J.L.D. designed the experiments and wrote the manuscript.

